# Statistical learning of large-scale genetic data: How to run a genome-wide association study of gene-expression data using the 1000 Genomes Project data

**DOI:** 10.1101/2022.09.03.506492

**Authors:** Anton Sugolov, Eric Emmenegger, Andrew D. Paterson, Lei Sun

**Author notes:** These authors contributed equally to this work (Anton Sugolov, Eric Emmenegger).

## Abstract

Teaching statistics through engaging applications to contemporary large-scale datasets is essential to attracting students to the field. To this end, we developed a hands-on, week-long workshop for senior high-school or junior undergraduate students, without prior knowledge in statistical genetics but with some basic knowledge in data science, to conduct their own genome-wide association studies (GWAS). The GWAS was performed for open source gene expression data, using publicly-available human genetics data. Assisted by a detailed instruction manual, students were able to obtain ∼1.4 million p-values from a real scientific study, within several days. This early motivation kept students engaged in learning the theories that support their results, including regression, data visualization, results interpretation, and large-scale multiple hypothesis testing. To further their learning motivation by emphasizing the personal connection to this type of data analysis, students were encouraged to make short presentations about how GWAS has provided insights into the genetic basis of diseases that are present in their friends and/or families. The appended open source, step-by-step instruction manual includes descriptions of the datasets used, the software needed, and results from the workshop. Additionally, scripts used in the workshop are archived on Zenodo to further enhance reproducible research and training.

## 1 Introduction

The overarching goal of this project is providing an example of engaging education in statistics to attract senior high-school or undergraduate students to the field, who will eventually grow and mature as competent data scientists. To achieve this goal, we designed a week-long workshop that provides students contextual, immersed, and hands-on learning experience in data science, using publicly-available, contemporary datasets.

We chose genetic data as the domain knowledge because they are complex, large-scale, high-dimensional, and practically important (Risch & Merikangas 1996). Although we do not expect nor want all students to continue their studies in statistical genetics, at the end of the workshop we expect students to (a) know about the variations in the human genome and the structure of the human population, (b) put into use their statistical knowledge by working with the 1000 Genomes Project (1KG) data (Auton et al. 2015), and (c) deepen their statistical understanding in areas including confounding (Cummiskey et al. 2020, Hu & Ziv 2008), heterogeneity (Higgins & Thompson 2002, Gordon et al. 2020), using principle component analysis to capture population structure (Price et al. 2006, Reich et al. 2008, Abdi & Williams 2010), multiple hypothesis testing (Shaffer 1995, Dudbridge & Gusnanto 2008), results interpretation and data visualization (Chanock et al. 2007, Hudiburgh & Garbinsky 2020), and reproducible research (Dragicevic et al. 2019, Ostblom & Timbers 2022).

In the last 15 years, genome-wide association studies (GWAS) have become a highly efficient way to identify genetic variants associated with traits and diseases (Manolio 2010, Visscher et al. 2017, Lappalainen & MacArthur 2021, Tan & Timpson 2022). The typical method involves testing millions of bi-allelic single nucleotide polymorphisms (SNPs), one-at-a-time for association with an outcome (e.g. the continuous blood pressure or the binary trait of high blood pressure) using either linear or logistic multivariate regression, and more recently generalized linear mixed-effect models (Cordell & Clayton 2005, Zhou & Stephens 2012). Although the commonly used statistical models are relatively simple for each SNP, the main challenge relates to the size of the human genome and the number of SNPs. For example, in imputed genetic data from the UK Biobank (Bycroft et al. 2018, Das et al. 2016), about 10 million SNPs are typically analyzed. Additionally, prior to association testing, several (domain-specific) quality control (QC) steps are necessary to restrict the analysis to SNPs and individuals with high quality data (Marees et al. 2018).

Most individual-level genome-wide SNP data is not publicly available due to privacy (Lunshof et al. 2008). We chose to illustrate GWAS using publicly available trait and genetic data from the 1000 Genomes project, in which participants consented to their data being made freely available (Clarke et al. 2012). Due to the small sample size available (about 1000 in total with 88 Yoruban and 102 Utah individuals, small relative to 1,344,840, the number of SNPs analyzed), we chose a trait that is known to be strongly associated with some SNPs with large genetic effects. This way, there would be sufficient power to detect the association with the small sample size; the remaining SNPs serve as negative controls and demonstrate issues pertinent to large-scale multiple hypothesis testing.

There is a wide variation in the level of gene expression in a specific tissue or cell, and much of this variation is influenced by SNPs near to a specific gene, the so-called cis-eQTL (Võsa et al. 2021). We used an example from earlier literature to illustrate the identification of genetic variation associated with the level of expression of the gene named Endoplasmic Reticulum Aminopeptidase 2 (*ERAP2*) (Cheung et al. 2005). The ERAP2 gene expression levels were measured in peripheral blood B cell lines in Utah residents with European ancestry, and Yoruba people from Ibadan, Nigeria from the 1000 Genomes Project (Roslin et al. 2016). The project is a publicly available catalogue of individual-level human genetic variation^1^, constructed by measuring genetic variation with an array of technologies in multiple populations around the world (Auton et al. 2015).

The workshop is designed to be executed with a 4-5 day period. The mornings can be used for the more traditional teaching modus operandi via (interactive) lectures, while the afternoons may be dedicated to the hands-on component with sufficient Teaching Assistant (TA) support. The student-TA ratio could range from 1-5 to 1-10, depending on the readiness of the student cohort. The last 2-3 hour of the workshop is recommended for a general discussion and obtaining feedback from the students, and ideally including short student presentations; see Section 2.5.

## 2 Methods

First and foremost, the workshop provides extensive hands-on experience in conducting, summarizing and interpreting a genome-wide association study to senior high-school students or junior undergraduate students with basic knowledge in data science. The hands-on experience includes using R (Maindonald 2008), running PLINK v1.9 (Purcell et al. n.d.) which is specific to the GWAS domain, and working with large-scale data. A detailed manual is attached as an Appendix, and the most updated version is openly accessible^2^.

Additionally, the workshop has the more traditional teaching/learning component through (interactive) lectures, covering complementary topics in genetics and statistics. We have made the lecture notes openly accessible^3^.

### 2.1 Datasets

In total, 190 individuals and 1,344,840 bi-allelic SNPs from the 1000 Genomes Project (Auton et al. 2015, Roslin et al. 2016) passing quality control from The Centre for Applied Genomic (TCAG)^4^ were used for the genome-wide association study.

Quality control is a significant component of conducting a proper GWAS (Marees et al. 2018). However, in-depth QC is domain-specific and time-consuming, not suitable for the purpose of this workshop. We thus provides a set of good quality data while emphasizing the importance of QC, so that the participating students could successfully carry out a preliminary GWAS within the first two days of the workshop and obtain *∼*1.4 million p-values from a real scientific study. We note that this early success is critical to keeping the students engaged and motivated to learn the theories that support their empirical results.

Cheung et al. (2005) identified that the expression of the gene *ERAP2* had strong genetic association in HapMap 3 individuals (International HapMap Consortium 2007), many of which overlapped with the 1KG individuals. Gene expressions of *ERAP2* measured in peripheral blood B cell lines were first extracted from Array Express (Montgomery et al. 2010, Stranger et al. 2012), then matched to the IDs of 1KG individuals, and finally formatted for PLINK; see Appendix A.

The two largest 1KG sub-populations are Yoruban individuals in Ibadan, Nigeria (YRI), and Utah residents (CEPH, Centre d’Etude du Polymorphisme Humain) with Northern and Western European ancestry (CEU). In total, 91 YRI individuals and 104 CEU individuals matched between the 1KG and HapMap 3 datasets, and they were used for the workshop purpose.

Using principal component analysis (PCA) of PLINK v1.9 (Purcell et al. n.d.), three and two outliers were removed, respectively from the YRI and CEU samples. Thus, the final GWAS analysis was restricted to 88 YRI individuals and 102 CEU individuals, and their genetic data of 1,344,840, bi-allelic SNPs. The basic PCA analysis pipeline is provided in the appended manual and could be part of the workshop if time permits.

### 2.2 Software

An introduction to PLINK (v1.90 beta 6.24) (Chang et al. 2015, Purcell & Chang 2021) is necessary for the purpose of this GWAS workshop. Depending on the readiness of the student cohort (and length of the workshop), a brief introduction to using R (v4.1.0) (R Core Team 2021) could be also part of the workshop; open-resource R introduction materials abound^5^.

PLINK is a command line toolkit for performing the GWAS computation efficiently, giving students hands-on experience with the most popular software used in the ongoing GWAS research. Additionally, the installation and use of R packages such as “qqman” (Turner 2018), “ggplot2” (Wickham 2016) and “hexbin” (Carr et al. 2021) introduce students to effective data visualization, a core component of interpreting GWAS results. Included in the open source manual is also a brief introduction to an (optional) use of the UNIX environment.

### 2.3 Overview of the Workshop Content

We summarize the main steps of running a GWAS of the gene expression data of *ERAP2*, using the 88 YRI individuals and their 1,344,840 SNP data of the 1000 Genomes Project (i.e. the YRI GWAS). We refer the readers to the open source manual and scripts, which include further analyses (i.e. the CEU GWAS) that could be reproduced using the step-by-step instructions.

Since trait distribution and SNP frequency may differ between populations, GWAS is often performed separately for each population (Price et al. 2006). In the analyzed sample, additional PCA may be conducted to capture fine-scale population structure (Reich et al. 2008); see Section 3 of the appended manual on population stratification.

#### 1. Prepare the datasets

Extract the cleaned 1KG SNP data into a separate analysis-specific directory.

First, students should specify the phenotype of interest and remove individuals who are not needed for the YRI GWAS. Students achieve these with the `--pheno` and `--prune` PLINK commands respectively; for additional details see the section named ‘Standard data input’ of the PLINK documentation^6^.

Second, students remove rare SNPs (e.g. with a minor allele frequency (MAF) less than 5%) and the sex chromosomes from the analysis using the `--maf 0.05` and `--chr 1-22` flags, respectively ^7^. (The 1000 Genomes data quality control performed by Roslin et al. (2016) does not include a MAF-based QC step.) Students should only analyze the autosomal common SNPs, as identifying associations on the sex chromosomes (Chen et al. 2021, Wang et al. 2022) and analyzing rare SNPs (Derkach et al. 2014) requires more intricate methods beyond the scope of the workshop.

Lastly, for computational reasons, students create binary files from this dataset with `--make-bed. The .bim, .bed, .fam` file types should be generated and named after `ERAP2_YRI`. Students should verify that the parameters they have entered are correct by viewing the `.log` file.

#### 2. Run the association analysis

Since gene expression data is continuous, students should specify a linear regression, with PLINK command `--linear`. This evaluates the association between the gene expression and each SNP, also known as the expression quantitative trait loci (eQTL) analysis.

Association analysis often includes covariates to avoid spurious associations from confounding. The sexes of the individuals are included in the dataset, so students may include this covariate in the eQTL GWAS analysis by using `--linear sex`.

#### 3. Post-association analysis and results interpretation

The association results can be sorted with `sort.R`, which also generates a file with the top 50 most significant SNPs.

The genome-wide results may be plotted and interpreted, which we explain with examples in the next section; also see the appended manual for additional details. Using appropriate QC steps, including the MAF filtering, prevents NA results in the output in principle. However, to be cautious the `NA_removal.R` script may be used to identify and remove NA results from the follow-up data visualization analyses.

### 2.4 A Highlight: Multiple Hypothesis Testing and Data Visualization

During the workshop, students are introduced to the multiple testing problem in GWAS through the morning lectures. Although the concept of multiple hypothesis testing, and its (theoretical) connection with ‘p-values being Unif(0,1) distributed under the null’, is covered in most introduction courses to statistics, student’s understanding and appreciation of this concept is often lacking, in part due to the traditional emphasis on identifying variables with p-values meeting some significance criterion, as opposed to exploring the whole distribution. This, in part, is a result from a lack of hands-on experience with large-scale real data analysis, With close to 1.4 million p-values obtained from a real GWAS, students realize that many SNPs (close to 70,000 in fact) are ‘significant’ if the traditional *α* = 0.05 type I error threshold were used. However, the histogram of p-values in Figure 1 shows an empirical distribution close to Unif(0,1), the distribution expected under the null hypothesis of no association. This is expected for a typical GWAS, as unless the trait is polygenic (i.e. with a large number of contributing SNPs) *and* the sample size is very large, most of the SNPs are not expected to be associated with the trait or their associations are not detectable (Devlin et al. 2001, Yang et al. 2011).

**Figure 1:**
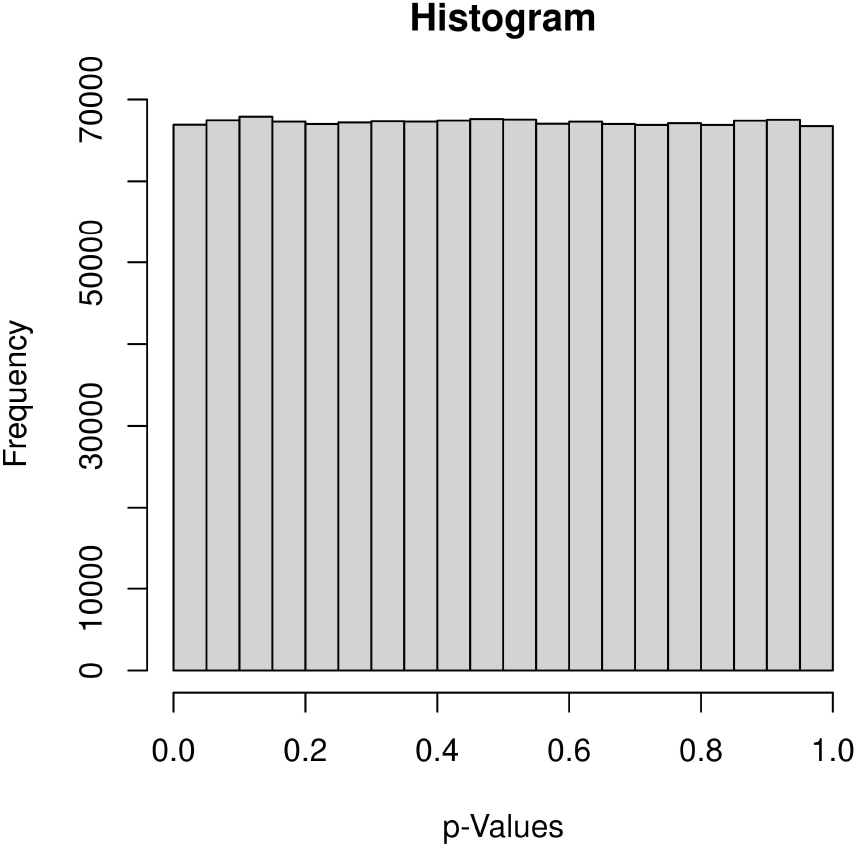
Histogram of the 1,344,840 p-values from the YRI GWAS of the gene expression of *ERAP2*, obtained using the workshop materials. The histogram is close to Unif(0, 1), the expected distribution of p-values under the null of no association.

Without going into the technical details, students are then introduced to the *α* = 5.0 *×* 10^*−*8^ genome-wide significance threshold used in GWAS to control the family-wise error rate at 0.05 (Dudbridge & Gusnanto 2008). Further, two most commonly used data visualization plots in GWAS are introduced: the Manhattan plot and the Q-Q plot as shown in Figure 2. These two plots complement the histogram which lumps all small p-values in one bin, thus masking the individual significant results.

**Figure 2:**
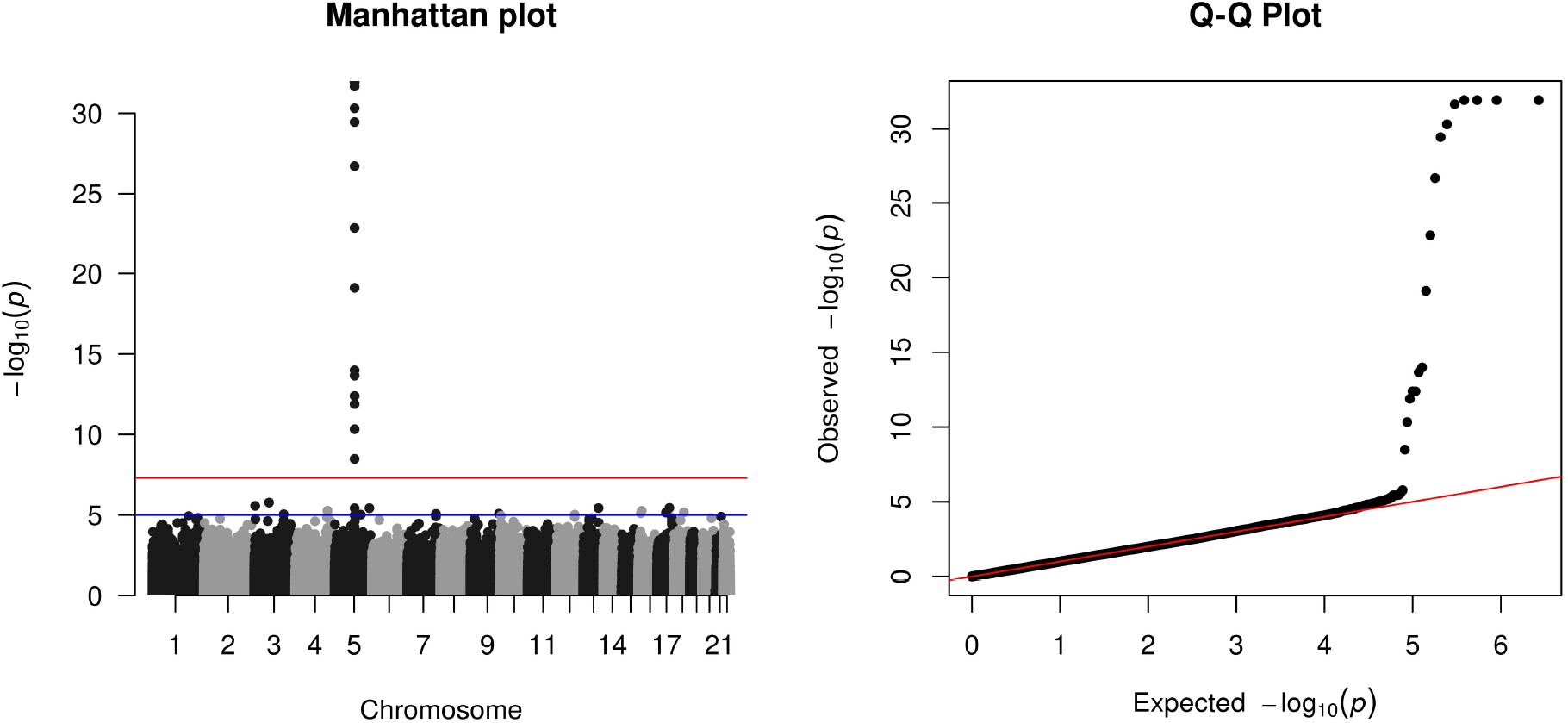
The Manhattan plot and Q-Q plot of the 1,344,840 p-values from the YRI GWAS of the gene expression of *ERAP2*, obtained using the workshop materials.

The Q-Q plot in Figure 2 is a standard statistical plot, showing the quantiles of the observed p-values against those of Unif(0,1), at the *−*log_10_ scale. In GWAS, the Q-Q plot serves two purposes. First, it highlights the significant results if there are any at the tail of the distribution. Second, it also shows the overall distribution of the GWAS p-values (though on the *−*log_10_ scale), which is typically expected to follow the main diagonal line.

Based on the Q-Q plot in Figure 2, it is clear that several SNPs are significantly associated with the gene expression of *ERAP2* in the YRI GWAS. However, their genomic locations (e.g. from which chromosome) are unclear. Thus comes the Manhattan plot which contrasts the *−*log_10_ p-value of each SNP against its genomic location, with the *α* = 5.0 *×* 10^*−*8^ genome-wide significance line (7.3 on *−*log_10_ scale) marked in red. Other significance thresholds for ‘suggestive’ association may also be shown, such as the *−*log_10_(10^*−*5^) blue horizontal line included in Figure 2.

In total, there are 17 genome-wide significant SNPs with p-values less than 5.0 *×* 10^*−*8^, all from the locus on chromosome 5 (at 96.2 – 96.3 Mb) that is close to the *ERAP2* gene. These are called cis-eQTL SNPs, i.e. SNPs near the gene and whose genotypes associated with differences in the gene expression level.

Another noticeable feature in a typical Manhattan plot is the ‘clustering’ of significantly associated SNPs. This is due to the phenomenon called linkage disequilibrium (LD) between nearby SNPs (Slatkin 2008). The location, p-values, and the LD between SNPs of a significant locus may be visualized in a Manhattan-like plot using the LocusZoom service (Boughton et al. 2021), with steps and the *ERAP2* example included in the appended workshop manual. Although LD is akin to the basic statistical concept of correlation, it is a more advanced concept in statistical genetics involving population genetics, thus not discussed further in this workshop.

### 2.5 Summary of the GWAS Workshop Conducted

In the summer of 2021, our team offered this workshop to 15 senior high school students from the University of Toronto Schools (UTS) in Toronto, Ontario, Canada. Due to the pandemic and limited number of TAs available, it was offered online and restricted to 15 participants, which were selected based on their interests and readiness in statistics, genetics and computing; see Appendix C for the application form. Post-workshop, a survey was conducted to collect participant feedback; see Appendix D for the survey questions.

Prior to the workshop, in addition to the survey, an earlier version of the appended manual was distributed to the participating students. Additionally, given the relatively low readiness of the participating students, the two lead TAs (AS and EE) provided detailed instructions for software installation and configuration, with a troubleshooting guide. Students followed this manual to work in groups, with clarification from the TAs via an online tutorial session as well as Discord discussions; Discord was the preferred social media of this group of students. At the time of the workshop, AS and EE were first year undergraduate students majored, respectively, in mathematics and life sciences, at the University of Toronto; AS and EE were mentored by ADP and LS during the summer of 2020.

Throughout the 4.5-day workshop, the morning lectures providing the necessary background in genetics and statistics were given, respectively, by ADP and LS. The afternoon sessions were guided tutorials, lead by AS and EE with participation of ADP and LS. Notably, on the last morning, students were encouraged to select a trait from the GWAS catalog^8^ (Buniello et al. 2019) and to present a 3-5 minute summary of a paper that performed a GWAS for that trait. In addition, students were encouraged to describe their motivation for selecting each particular trait, which provided an emotional connection to the science through personal stories, typically related to family history of diseases. The presented traits ranged from gout, breast cancer, to multiple sclerosis. Finally, to keep the students engaged, music were curated in advance and played during the (frequent) breaks, and the song “Another Brick in the Wall”, by Pink Floyd, was much appreciated by the students based on their feedback.

## 3 Student Feedback

After the workshop, a feedback survey (Appendix D) was distributed and eight responses were collected. Students found the workshop overall interesting, especially working with and interpreting the genetic component of the workshop. The students particularly enjoyed the SNP finding activity, and found the guided afternoon sessions helpful to their understanding.

Due to the high school background of the students, and the workshop’s limited time frame, some found the pace of the lectures to be overwhelming, particularly the statistical section of the lectures. Students unaccustomed to programming found using the terminal-based PLINK to be confusing, and recommended adding a terminal tutorial to the workshop manual, which was later included.

## 4 Discussion

Depending on the experience of participants, the scope of the workshop may be extended, including covering more advanced lectures, analyses and plots, as well as analyzing additional datasets. Discussions around the cleaning of 1000 Genomes data could be included in the morning lecture sessions, and cleaning steps for 1000 Genomes individuals (Roslin et al. 2016) may be replicated in the afternoons. More thorough descriptions of large-scale multiple testing and fundamentals of regression in the GWAS context may be included. An analysis using individuals with different populations, with PCA adjustment, may be given in the practical hands-on sessions. After conducting a sample GWAS in one population (e.g. the YRI GWAS), gene expressions with various significance (Cheung et al. 2005) matched with other 1KG populations may be provided for students to replicate. Included UNIX commands may be used as an introduction to conducting a remote GWAS on a cloud-based system, which typically are typically UNIX-based.

To adhere to the current standard of reproducible research (Peng 2011), initial GWAS were conducted and documented independently by AS and EE. The two sets of results were then compared with each other, and the analyses and results were successfully reproduced, independently, by the workshop participants. Additionally, the observed *ERAP2* significance replicates the earlier work by Cheung et al. (2005). R, PLINK, and dataset versions were synchronized, and all scripts were version-controlled and hosted on GitHub^9^. The exact analytical steps were recorded in a GWAS documentation, which would later become the appended, open source manual that allows users to reproduce the workshop GWAS materials. Finally, the tested workshop datasets and other materials were made publicly available on Zenodo ^10^.

## Supporting information

Workshop Instruction Manual

## 5 Acknowledgement

This research is funded by the Natural Sciences and Engineering Research Council of Canada (NSERC, RGPIN-04934), the Canadian Institutes of Health Research (CIHR, PJT-180460), and the University of Toronto Data Sciences Institute (DSI) Catalyst Grant.

## 6 Conflict of Interest Statement

The authors declare that the research was conducted in the absence of any commercial or financial relationships that could be construed as a potential conflict of interest.

## APPENDICES

### A Phenotype Extraction and Dataset Generation

Please refer to Github.com/sugolov/GWAS-Workshop/Notebooks/DatasetPreparation.Rmd to create a phenotype file using the expression data from the University of Geneva Medical School Montgomery et al. (2010), Stranger et al. (2012). Example phenotype files for ERAP2 are provided for the CEU and YRI populations both separately and together at Github.com/sugolov/GWAS-Workshop/Datasets. Please refer to Github.com/sugolov/GWAS-Workshop/Notebooks/YRI_Analysis.Rmd to combine the phenotype files with the 1KG dataset from The Centre for Applied Genomics Auton et al. (2015), Roslin et al. (2016)

### B High Coverage Dataset

The High Coverage dataset was generated using the 30x High Coverage samples from the New York Genome Center (NYGC) Byrska-Bishop et al. (2021). Please refer to Github.com/sugolov/GWAS-Workshop/Notebooks/ High_Coverage.Rmd to generate a set of High Coverage data.

### C Application Form

The application form for students consisted of the following questions sent out as a Google Form.

1. Your name (First, Last)
2. Your email address
3. Please list relevant courses (UTS course codes and names) taken in statistics/data science, computer science and biology. (This is to help the workshop organizers to team up participants with complementing skills, if needed depending on the number of applicants.)
4. Check 1-2 boxes that reflect your strengths
  - Statistics/Data Science
  - Computing
  - Biology
5. Explain why are you particularly interested in this workshop? (200 words)
6. Any preference or suggestion on the platform(s) to be used for the virtual workshop, and for the on-line discussion board?
7. Any other comments?

### D End of Workshop Survey

The following questions were sent to the students as a Google Form after the end of the workshop. 8 students out of 17 responded.

1. On a scale of 1 to 10, how difficult did you find the genetic component?
2. If you answered greater than 7 to the question above please specify what you found too difficult. If you answered below 5 to the question please specify what you found too easy. If <=5 your answer <=7, still say something:-)
3. Did you find the pace of the genetic component too quick or too slow? Please specify.
4. What would you have liked to seen more of?
5. On a scale of 1 to 10, how difficult did you find the statistics component?
6. If you answered greater than 7 to the question above please specify what you found too difficult. If you answered below 5 to the question please specify what you found too easy. If <=5 your answer <=7, still say something:-)
7. Did you find the pace of the statistic component too quick or too slow? Please specify.
8. What would you have liked to seen more of?
9. On a scale of 1 to 10, how difficult did you find the computing/hands on component?
10. If you answered greater than 7 to the question above please specify what you found too difficult. If you answered below 5 to the question please specify what you found too easy. If <=5 your answer <=7, still say something:-)
11. Did you find the pace of the computing component too quick or too slow? Please specify.
12. What would you have liked to seen more of?
13. If we were to do this workshop again what would you have liked to see more of? Select all that apply.
  - Statistics
  - Genetics
  - Computing
  - Nothing. The balance was perfect.
  - Other:
14. What was your favourite aspect of the workshop? Select all that apply.
  - Statistics
  - Genetics
  - Computing
  - None. I did not enjoy anything
  - All. I loved everything
  - Other:
15. Would you like to have been presented with more references and resources before the workshop (ie. terminal commands, file directory structure, etc)? If this is the case please specify.
16. From a scale of 1 - 10 how much did you enjoy the music during breaks?
17. Any other final remarks?

## Footnotes

1 https://www.internationalgenome.org/about

2 https://github.com/sugolov/GWAS-Workshop

3 https://github.com/LeiSunUofT/How-to-Run-a-GWAS

4 https://tcag.ca/tools/1000genomes.html;https://www.internationalgenome.org/

5 https://cran.r-project.org/doc/manuals/r-release/R-intro.pdf

6 https://www.cog-genomics.org/plink/

7 https://www.cog-genomics.org/plink/1.9/filter

8 https://www.ebi.ac.uk/gwas/

9 https://github.com/sugolov/GWAS-Workshop/

10 https://zenodo.org/record/6981694

## Notes

https://zenodo.org/record/6981694

### Competing Interest Statement

The authors have declared no competing interest.

