## Supplementary material for "Statistical learning of large-scale genetic data: How to run a genome-wide association study of gene-expression data using the 1000 Genomes Project data": Workshop Instruction Manual

### Genome-Wide Association Study Workshop Manual

Anton Sugolov and Eric Emmenegger
*Supervised by Dr. Andrew Paterson and Dr. Lei Sun*

This guide is intended for participants in the UTS Genome-wide Association Workshop in June 2021. The guide includes an overview of the studies we will perform, as well as steps to install the necessary software. Please complete the steps in sections 1.2 - 1.3 before our first session or arrive ready to troubleshoot. We hope this manual will be an adequate reference.

**2 Introduction to Terminal Commands** **9**

**6 Conclusions** **34**

**7 Resources** **35**

### 0. Abbreviations

- **ERAP2:** Endoplasmic Reticulum Aminopeptidase 2
- **GWAS:** Genome-wide Association Study
- **PCA:** Principal component analysis
- **SNP:** Single Nucleotide Polymorphism

### 1. Preliminaries

#### 1.1. Introduction

In this workshop, we will learn how to perform a genome wide association study (GWAS). A GWAS tests the association between single nucleotide polymorphisms (SNPs) throughout the genome and a desired phenotypic trait. For example, given 100 individuals and their heights, we may set the phenotype as their height and attempt to identify what SNPs are most statistically significantly associated with it. This type of study has become popular due to the ability to identify associations with previously unknown risk factors. For example, given a new association between a genetic variant (SNP) and cardio-vascular disease, further studies may be performed about the mechanism for how this SNPs alters specific genes which then influence the onset of the illness. All published associations can be found within the NHGRI GWAS Catalogue (Buniello *et al.* 2019), including a visualization of their locations on chromosomes.[1].

In practice, running a proper analysis is not so simple. Suppose we conduct a GWAS on the gene expression of LCT (lactase) with participants that have differences in age and ethnic background. Due to different populations having different rates of lactose intolerances, due to different expression of the LCT gene, we may experience spurious associations caused by population differences. Similarly, lactose intolerance increases with age, which may cause more confounding. Since we aim to identify regions that are associated with lactase, and not regions that could give confounding associations between participants, we must account for covariates present within our studies. Considerations like these must be carefully treated in order for a GWAS to produce reliable results.

Additionally, trait heritability is another challenge to a proper analysis. Heritability is defined as how much of the variation of a trait is due to genetic variation, and thus how likely it is that the same trait will be inherited in the genes of offspring (Wray & Visscher 2008). Using the aforementioned example, height varies not just due to genetic factors, but also due to environmental factors such as nutrition. As these factors are typically extremely difficult to quantify, we will assume that only the genotype of an individual affects their trait, and that any other factors do not have any effect.

In this manual, we conduct an association analysis between the level of gene expression of *Endoplasmic Reticulum Aminopeptidase 2* (Cheung *et al.* 2005) measured in peripheral blood B cell lines separately in Utah residents with European ancestry, and Yoruba people from Ibidan, Nigeria from the 1000 Genomes Project (1KG) (Roslin *et al.* 2016). The 1000 Genomes Project is a catalogue of human genetic variation, constructed by measuring the genetic variation of various populations around the world Auton *et al.* 2015 [2]. Typically much rigorous quality control occurs before the genetic data can be used for a GWAS analysis. In the interest of time, we will use cleaned data provided by Dr. Andrew Paterson (Roslin *et al.* 2016). The gene expression data for the individuals we will be studying were taken from publicly available Illumina gene expression microarray data (Montgomery *et al.* 2010) (Stranger *et al.* 2012) [3] on 800 HapMap 3 individuals (International HapMap Consortium 2007), with significant overlap between 1KG individuals. We will use this data to identify the region(s) of the genome that are most significantly associated with ERAP2 gene expression, that is close to the gene encoding ERAP2 itself.

[1] https://www.ebi.ac.uk/gwas/diagram
[2] https://www.internationalgenome.org/about
[3] https://www.internationalgenome.org/category/gene-expression/

#### 1.2. Required Software and Data

We will mainly be using R and PLINK v1.9 to conduct our analyses.

R v4.1.0 is a language useful for various statistical computations and data visualization purposes. It will mainly be used to interpret the results of our GWAS. R is free, has an easy to learn syntax, and provides a great environment for installing packages for tailored purposes. A brief summary about R installation on Windows and MacOS will be provided in section 1.3. In addition to R, we recommend using R-studio as the Integrated Desktop Environment (IDE) to perform the steps in this manual. For more information, the R website is linked below.

`https://www.r-project.org/`

There are a variety of resources helpful for learning R. The documentation to any package in R can be found here:

`https://www.rdocumentation.org/`

Syntax and documentation for packages can be found above. A textbook that provides a solid foundation for using R is *Hands-On Programming with R* written by Garrett Grolemund (Grolemund 2014). The first chapter gives an overview of the basic ideas in R, and will be sufficient for the purposes of this workshop. Since there is no online access to this textbook through the University of Toronto Library, this resource is not mandatory, but is recommended if it can be found. Further guides to using R (Maindonald 2008) can be found below:

`https://www.rdocumentation.org/`
`https://cran.r-project.org/doc/contrib/usingR.pdf`

The above is somewhat dated, but is relevant for the basic ideas. For specific questions about R, it is better to search for relevant posts in the StackExchange as they are often most useful.

PLINK v1.90 is a free and open-source command-line program for performing genome wide association studies. It was originally developed in 2007 by Shaun Purcell (Purcell *et al.* 2007) with version name v1.07, but was recently updated and released as version v1.90 by Christopher Chang (Chang *et al.* 2015). PLINK offers a simple way to work with the files in a GWA study. The features most necessary for our purposes are data management and association testing. If you have little experience with using the terminal in order to perform tasks, working in this way may require some learning. Fortunately, the syntax of PLINK is also easy to learn and provides a simple way to perform powerful analyses. The installation of PLINK is more involved, and a guide is included in 1.3. The documentation for all PLINK commands and instructions can be found at

`https://www.cog-genomics.org/plink/`

The above documentation for PLINK v1.90 is the most accurate reference for commands. To understand the syntax of the options for each function, read the "Interpreting our flag usage summaries" section on the following part of their site:

`https://www.cog-genomics.org/plink/1.9/general_usage`

Some portions of PLINK syntax and file structure have stayed the same since v1.07. The documentation to the original version of PLINK is also often useful, and is linked below:

`http://zzz.bwh.harvard.edu/plink/`

At this site there are tutorials and information about basic usage and data formats. **Please read the the first half of the section about PED files, first half about MAP files, all of binary PED files, and first third about alternate phenotype files at this link:**

`http://zzz.bwh.harvard.edu/plink/data.shtml`

**Note:** Appropriate UNIX commands will be included throughout this manual since they simplify some steps.

#### 1.3. Setup and Installation

##### 1.3.1. R - Windows

Downloading R for Windows is straightforward and is done through an installation wizard. The installer for the most recent version of R can be downloaded here:

`https://cran.r-project.org/bin/windows/base/`

Once you have downloaded it, run the installer and proceed with the installation. **Ensure that the installation adds R to your system PATH.** Check that R is installed by successfully running `R` in your Command Line.

##### 1.3.2. R - MacOS

To install R on MacOS, the first step is to go to the following link, clikck on "Download for (Mac) OS X" and download the most recent pkg file.

`https://cloud.r-project.org/`

Locate the pkg file (default is in the Downloads directory). Optionally, you can check the integrity of the file using either of the commands (Sha1 checksum or pkgutil) provided. Double click on the file to install R. Only the base package is required, so you do not need to install some of the options such as Texinfo. Check that R is installed by successfully running `R` in your Terminal.

##### 1.3.3. R Packages - MacOS and Windows

In order to generate some of the plots the R packages `qqman`, `hexbin` and `ggplot2` will be necessary. To install these, open R in your terminal, and run the following commands

```
install.packages("qqman", repos = "https://cran.r-project.org/")
install.packages("ggplot2", repos = "https://cran.r-project.org/")
install.packages("hexbin", repos= "https://cran.r-project.org/")
```

**Note:** You may need to be signed in as an administrator/super user to run these.

##### 1.3.4. PLINK - Windows

First go to

`https://www.cog-genomics.org/plink/`

Download the latest stable release for Windows and your system architecture [4] (x86/32-bit or x64/64 bit) given in the table. Once downloaded, extract the .zip file into a new directory. `plink.exe` should be one of the extracted files. You can now click and run `PLINK`. To make `plink` available in any directory, the executable which is most easily done by adding it to a directory in your system path. This can be done by copying `plink.exe` into

`C:\Program Files`

You can navigate to the above directory and simply drag `plink.exe` into it. Otherwise, use the Command Prompt to navigate to the directory where `plink.exe` is located and run

`cp plink.exe C:\Program Files`

Check that PLINK is installed by successfully running `plink` in your Command Line
**Note:** You will need to be the system administrator to do the above.

[4] You can check your system architecture in `Settings > About > System Type |`

##### 1.3.5. PLINK - MacOS

To install Plink on MacOS, please first go to

`https://www.cog-genomics.org/plink/`

Download the latest stable release given in the table. Double click on the .zip file to extract the archive. If you open this directory in a Terminal, you can try some of the PLINK commands on the toy file set. To be able to use PLINK in any directory, you will need to copy the binary into a directory in your system PATH. To do so, we will first make a directory, then add it to the path, and then add Plink to that directory. Open a Terminal, and copy in each line below one at a time (for the last line, make sure you are in the directory which contains the PLINK binary).

```
mkdir ~/bin
export PATH=$PATH:~/bin/
cp plink ~/bin
```

Now, you can run Plink in any directory. Check that PLINK is installed by successfully running `plink` in your Terminal.

##### 1.3.6. Optional - Homebrew installation for MacOS

If opting to use specific Unix commands, it will be necessary to install the GNU's Not Unix! (GNU) core utilities. Specifically, MacOS comes with Berkeley Software Distribution (BSD) `grep`, but GNU `grep` contains a flag that may be necessary. The easiest way to install GNU `grep` is through Homebrew. Homebrew is a package manager which can be used to install other free software as well. To install Homebrew, navigate to:

`https://brew.sh/`

Copy the command given under the Install section of the website into a Terminal, and press enter. Once Homebrew is installed, install the `coreutils` package by entering the following command into a Terminal

```
brew install coreutils
```

##### 1.3.7. Data

The cleaned 1KG data will be provided by Dr. Andrew Paterson and his team at The Centre for Applied Genomics. **Download the PLINK files for the independent samples here:**

`http://tcag.ca/tools/1000genomes.html`

**Required 1KG files**

- PLINK binary format files of independent samples:
  `295.19 MB indep.tar.gz`

Note the quality control steps performed on the original 1KG data, and the various other statistics included in the tables. The original ERAP2 gene expression data from the Illumina Human-6 v2 array is available here:

`https://www.internationalgenome.org/category/gene-expression/`

and the annotation file for the gene names is available here:

`https://www.ncbi.nlm.nih.gov/geo/query/acc.cgi?acc=GPL6102`

Extracting the data for ERAP2 is time consuming, and so we've already extracted the files we need for this analysis. **Download the files available here**:

`https://github.com/sugolov/GWAS_Workshop`

**Required GitHub Files**

- ERAP2 Gene Expression for 102 CEU individuals:
  `41.1 KB, Datasets/ERAP2_CEU_phenotypes.txt`
- ERAP2 Gene Expression for 88 YRI individuals:
  `41 KB, Datasets/ERAP2_YRI_phenotypes.txt`
- ERAP2 Gene Expression for both 102 CEU and 88 YRI individuals:
  `42.4 KB, Datasets/ERAP2_CEU_YRI_phenotypes.txt`
- Directory containing relevant R scripts for plotting:
  `14.8 KB, Scripts`

### 2. Introduction to Terminal Commands

As mentioned in Section 1.2, Plink is operated through a command-line interface (CLI). This means that it does not have a traditional graphical user interface and that it must be used through the Terminal on MacOS and Linux, or the Command Prompt on Windows. Accordingly, it is helpful to learn some basic operations in a CLI. For example, if you open the file browser on your computer (Finder on MacOS, File Explorer on Windows), you are greeted by your folders and files. Folders can also be referred to as directories. You can use your mouse to click on both, and perform operations on such as opening, renaming and deleting. You can do all of these operations in the command line as well.

#### 2.1. MacOS and Linux Terminal Commands

Table 2.1 shows basic commands that can be used in MacOS, and Linux, which are both Unix-like operating systems. Note that each command also has different options. For example, while the command `rmdir` can only be used for deleting empty directories, the command `rm -r` can be used to delete directories and all their contents. To learn more about all options, use the `man` command.

Table 1: Basic Terminal Operations and their corresponding commands in Unix-Like Operating Systems

| Operation | MacOS/Linux Command |
|---|---|
| Copy a file from one location to another | `cp [SOURCE] ... [DESTINATION]` |
| Move a file from one location to another | `mv [SOURCE] ... [DESTINATION]` |
| Delete a file | `rm [FILE]` |
| Change directories | `cd [TARGET DIRECTORY]` |
| List the contents of your current directory | `ls` |
| Make a new sub-directory in the current directory | `mkdir [DIRECTORY NAME]` |
| Remove an empty sub-directory in the current directory | `rmdir [TARGET DIRECTORY]` |
| Show the help page for a command | `man [COMMAND NAME]` |

#### 2.2. Windows Command Line Commands

Table 2.2 shows basic commands for Windows.

Table 2: Basic Terminal Operations and their corresponding commands in Unix-Like Operating Systems

| Operation | Windows Command |
|---|---|
| Copy a file from one location to another | `cp [SOURCE] ... [DESTINATION]` |
| Move a file from one location to another | `mv [SOURCE] ... [DESTINATION]` |
| Delete a file | `rm [FILE]` |
| Change directories | `cd [TARGET DIRECTORY]` |
| List the contents of your current directory | `dir` |
| Make a new sub-directory in the current directory | `mkdir` or `md [DIRECTORY NAME]` |
| Remove a sub-directory in the current directory | `rmdir [TARGET DIRECTORY]` |

#### 2.3. Tips and Tricks

Note that examples will be provided with Unix commands only, refer to Tables 2.1 and 2.2 to translate commands to Windows.

- **Tab completion:** Entering the first few letters of a file or folder and pressing Tab will autocomplete its name. If you type in the first few letters, and they are not unique to one folder or file, then all possible options will show when you press Tab.

  **Example:** If you are in a directory containing two sub-directories called Documents and Downloads, and you type in `cd Do` and press Tab, then Documents and Downloads will appear below. As soon as you type in either a c or w, there is only one directory which fits the pattern Doc or Dow, so pressing Tab will autocomplete the full directory name. That is, if you type in `cd Doc` and press Tab, it will autocomplete to `cd Documents/`.

- **Using the * character:** The character * can be used to signify all files and sub-directories.

  **Example:** The command `mv * [TARGET DIRECTORY]` will move all files and sub-directories in the current directory to the target directory.

- **Using the up arrow:** In command-line interfaces, the up arrow can be used to access the last commands you ran. That is, if you press the up arrow once, it will show the last command you ran; if you press the up arrow twice, it will show the second last command you ran, etc. This can be useful if you are running the same command but with different files.

  **Example:** Say the last command I ran was:

  ```
  plink --bfile ERAP2_YRI --linear --out ERAP2_YRI_sex
  ```

  If I press the up arrow, then this will appear, and I can make changes to the command before running it again.

### 3. Quality Control

The genetic data used for an analysis must undergo quality control so that the GWAS gives reliable results (Marees *et al.* 2018). Poor quality of DNA samples, genotype probes, or sample contamination can seriously influence the reliability of the analysis. Approximately 90% of the time spent performing a GWAS is dedicated to quality control. Given the time and scope of this workshop, we will skip performing the majority of these steps and instead describe them. We will perform principal components analysis on the populations as a means of quantifying variation between populations in our later analyses. The data we will be using has already been cleaned by Dr. Andrew Paterson's team[5]. In addition to the general steps provided by Marees *et al.* (ibid.), we will include the specific thresholds used in the quality control of our data.

1. **Missingness of SNPs and Individuals.** Individuals that have a large genotype missingness or SNPs that that are missing in a large proportion of the subjects are removed. This step is included to ensure general sample quality.
   - The average proportion of non-missing genotypes was very high in the sample at 98.3%. Individuals below the commonly accepted 97.0% threshold were excluded.
2. **Sex Discrepancy.** Checks for discrepancies between the reported sex of individuals and their sex chromosome composition. A large number of discrepancies can indicate sample mixups. In some cases the sex can be inferred from chromosome composition.
   - Inferred sex was consistent with provided sex for 99% of individuals, and the sex of two individuals was imputed from their genotypic data.
3. **Minor Allele Frequency.** SNPs with a low minor allele frequency lack power in detecting associations, and are more prone to genotyping errors. These are typically excluded based on empirical thresholds.
4. **Heterozygosity.** Heterozygosity is a term for an individual having different alleles for a SNP. This step excludes individuals that deviate from the mean rate of heterozygosity in the sample, as this can indicate sample contamination or inbreeding.
   - The average autosomal heterozygosity was calculated to be 0.20, with no unusually extreme samples. The wide distribution (figure 1) of heterozygosities is consistent with the multi-ethnic nature of individuals from the 1KG project.
   - Individuals with an unusually high number of heterozygous genotypes in the mitochondrion, male X chromosome, and Y chromosome could provide evidence of quality issues. None were removed as they were not detected in the data.
5. **Relatedness.** Relatedness of individuals above a threshold can produce false positive association due to non-independence, and are removed. Related individuals are sometimes not removed but instead taken into account using special methods when performing a GWAS on a family based sample.
   - Complete pedigrees were constructed using this data for dummy individuals, but will not be used in our analyses.

[5]Please see `http://tcag.ca/documents/tools/omni25_qcReport.pdf`.

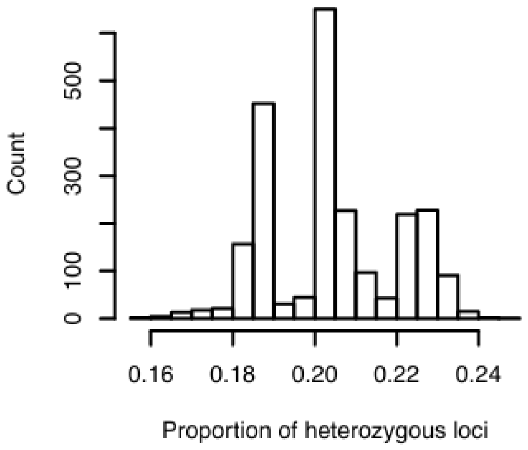

Figure 1: Histogram of proportion in heterozygous loci from data cleaned by Roslin *et al.* (2016).

6. **Population stratification.** We will perform this through principal component analysis (PCA). This process generates a set of data that quantifies the relationships between ancestral sub-groups in the sample. This data is then included as a covariate in our analysis, or analysis is performed taking into account the genetic differences between different populations. This step is crucial as population stratification is an important source of systemic bias in GWAS.

In total, 1,317,318 high quality SNPs passed the quality control measures (Roslin *et al.* 2016). These SNPs have an average proportion of non-missing genotypes >97%, two observed alleles, and no errors in Mendelian transmission.

We urge you to take a look at some of the quality control measures by Roslin *et al.* (ibid.) conducted on the data we will be using for our analysis, and from which the above summary statistics are taken. This will give you some additional insight to one of the most critical stages of conducting a GWAS, and how to perform it. We also recommend that you read the A. Marees paper *A tutorial on conducting GWAS: Quality control and statistical analysis* where this sequence of steps was taken from. It provides an outline of how to use PLINK in addition to various other useful information about conducting GWAS. If you are interested, you should perform the QC tutorial on his GitHub (`https://github.com/MareesAT/GWA_tutorial/`) as it is very instructive.

After quality control is complete, the GWAS can begin.

### 4. Analysis

First, a GWAS on 88 YRI individuals will be performed. 102 CEU individuals and their gene expression will be provided afterwards, and you will be responsible for performing an independent GWAS for these individuals. Annotated R commands for visualization will be included after each step.

You will often use R to read a data table. The `read.data.table()` R function may be used in the following way

```
#reads a data table
#
#   header=TRUE            - reads the top row as names for the columns
#   sep = ""               - reads any length whitespace between text as separator
#   fill = TRUE            - matches row length with NA values where applicable
#   quote = ""             - indicates no data is
#   check.names = false    - does not check package specific naming conventions
#
#   https://www.rdocumentation.org/packages/utils/versions/3.6.2/topics/read.table

results <- read.table("data.txt",
                      header = TRUE,
                      sep = "",
                      fill = TRUE,
                      quote = "",
                      check.names = FALSE)
```

#### 4.1. Before an analysis

##### 4.1.1. Exploratory analysis of the phenotype

Before a GWAS is conducted, it is helpful to understand the trait being studied. Some simple plots can help greatly in understanding the data.

1. **Summary statistics**. Creating summary statistics quantifies many valuable aspects of the data. The counts, quartiles, mean, median, mode, standard deviation, minimum, and maximum help with values needed to generate other plots. These should be communicated in a presentation of results. Please see the following R commands.

```
#   x                      - an arbitrary vector with numerical values, for example
x <- as.vector(data)
avg <- mean(x)         #   - finds the mean of the values in x
med <- median(x)       #   - finds the median of the values in x
sd <- sd(x)            #   - finds the standard deviation of the values in x
var <- var(x)          #   - finds the variance of the values in x
min <- min(x)          #   - finds the minimum of the values in x
max <- max(x)          #   - finds the maximum of the values in x
```

2. **Phenotype histogram.** This plot gives visualizes the distribution of the trait being studied. The variance, and presence of outliers become apparent. Note that this only applies to continuous traits. For binary traits, knowing the counts of affected/unaffected individuals is most important. Histograms can be generated using the `hist()` command in R.

```
#generates the histogram
#
#   as.numeric(data)    - the data being plotted, numeric type of third column in the
#                           data table (gene expression is in the third
#                           column in the phenotype files)
#   main =                - the main title
#   xlab =                - the x axis title
#   ylab =                - the y axis title
#   breaks=seq(8,14,0.5)  - plots bins of width 0.5 taking values from 8 to 14
#   freq = TRUE           - plots frequencies
#   cex = 3               - increases scaling for better resolution

        hist(as.numeric(data),
             main="Gene Expression distribution",
             xlab="Gene Expression value",
             ylab="Frequency",
             breaks = seq(8,14,0.5),
             freq = TRUE, cex = 3 )
```

The histograms and summary statistics of the ERAP2 gene expression should be generated for each set of individuals being studied. Generating gene expression distributions for a sample of individuals from different populations is helpful in identifying whether the distribution and variance changes based on population. Note significant modality apparent in the data.

##### 4.1.2. Exploratory analysis of the individuals' genetic data

Histograms are useful for most data. They can be used to plat many statistics such as minor allele frequency and heterozygosity within the populations being studied. For example, in the quality control steps used to prepare the data we will be using (Roslin *et al.* 2016), it was noted that the heterozygosity histrogram of the individuals across all populations was trimodal:

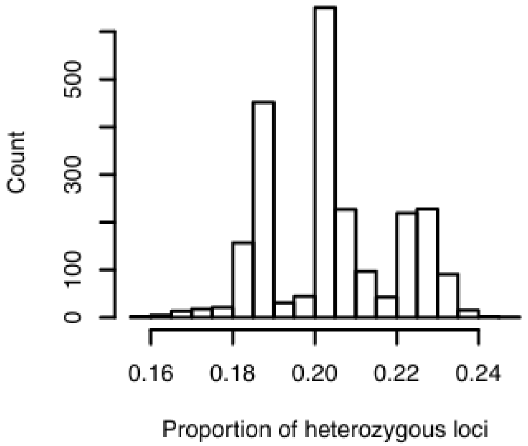

Figure 2: Histogram of proportion in heterozygous loci from data cleaned by Roslin *et al.* (2016).

The three modes present in the plot likely represent the world's major ethnic groups (ibid.), demonstrating that heterozygosity is a major statistic that varies significantly due to population differences. To gain an understanding of other metrics, we may generate histograms for descriptive statistics like the minor allele frequency. The minor allele frequency (MAF) is the frequency at which the second most common allele of a SNP occurs. To retrieve the minor allele frequency per SNP in a dataset, the PLINK command `--freq` may be used:

```
plink --file data --freq --out data_maf
```

Other descriptive statistics like SNP per chromosome may also be generated in order to explore the genotypic data. Please see the PLINK documentation (section 1.3.5) for other useful genotypic descriptive statistics.

##### 4.2. ERAP2 gene expression GWAS on 88 YRI individuals

In the first analysis, we perform a complete GWAS on 88 YRI individuals from the 1000 Genomes project. From the HAPMAP_3 gene expression dataset (Stranger *et al.* 2012), 91 YRI individuals overlap with the data cleaned by TCAG (Roslin *et al.* 2016). After examining their principal components, 3 outliers were identified and removed. A GWAS on these 88 individuals and their ERAP2 expression will be outlined.

1. Extract the cleaned 1KG data into directory containing the YRI phenotype .txt file. First, we specify the phenotype and remove all individuals in the cleaned data that we will not be studying. We do the first by `--pheno` and the next with `--prune` [6] in PLINK. Then, create a new binary file from this dataset with `--make-bed`.

   ```
   plink --bfile indep --pheno ERAP2_YRI_phenotypes.txt --prune --make-bed --out ERAP2_YRI
   ```

   `.bim`, `.bed`, `.fam` type files should be generated and named after `ERAP2_YRI`. You should also always verify that the parameters you have entered are correct by viewing the `.log` file.

2. In order to eliminate possible confounding in individuals from different populations, we typically include population covariates in the association test (see Section 3 on Population Stratification). In this analysis we are dealing with a single ancestral population, so the step is unnecessary. The inclusion of principal components as covariates will become relevant in the third analysis, where a GWAS will be performed on both CEU and YRI populations.

3. We must prepare our dataset before we can perform the association tests. In our analysis, we will exclude SNPs with minor alleles that have a frequency less than 5% of the total allele pool (each individual carries two alleles, so the total allele pool is twice the population size) using the command `--maf 0.05`. SNPs with a low minor allele frequency will have too few individuals in the 'effective group' to give reliable p-values. We will only include the autosomes (non-sex chromosomes) in our analysis using the command `--chr 1-22`. Associations on sex chromosomes are more complicated and are outside the scope of this workshop. [7]

   ```
   plink --bfile ERAP2_YRI --chr 1-22 --maf 0.05 --make-bed --out ERAP2_YRI_clean
   ```

4. Since the gene expression data we are using is continuous, we may run a linear regression with covariates through `--linear`. The sex of the individuals is included in the dataset, so we simply specify to PLINK to use sex as a covariate during the test with `sex`. For the purposes of this analysis, we are not interested in the statistical significance of each covariate, so we hide it in our output file with `hide-covar`.

[6] Please see the links to the official PLINK documentation in section 7.

[7] The differences between the X and Y chromosomes results in complications with the comparison, especially given that sexes typically have different combinations of these chromosomes.

```
plink --bfile ERAP2_YRI_clean --linear sex hide-covar --out ERAP2_YRI_sex
```

The output should be a `.assoc.linear` file. You should verify these steps by viewing the `.log` file.

5. After these steps, you can remove NA results by modifying the script NA_`removal.R`. As well, the data can be sorted with `sort.R` which also generates a file with the top 50 most significant SNPs in the genome. After cleaning the results, they must be summarized to others.

**Note:** For sorting and NA value removal, UNIX users can use the following:

```
awk '!/'NA'/' input.file | sort -gk 9 > output.file
```

These can also be done separately, which may be necessary in some cases. In this case, separate the commands, and remove the pipe symbol "|" in between them.

```
awk '!/'NA'/' input.file > output1.file
sort -gk 9 ouptut1.file > output2.file
```

After completing these analyses, the results must be visualised. A list of several plots and text files is provided below, and will greatly help in communicating the results. Please see section 5 for more details.

##### 4.2.1. Output text files

1. **README.txt** A file containing a description of other files included in the directory, and a *brief* [8] summary of the analysis performed.
2. **PLINK .log files** Someone viewing these results may be interested in understanding what PLINK commands were used.
3. **List of top 50 SNPs** A file containing a list sorted by p-value of the top 50 most statistically significant SNPs in the results.

##### 4.2.2. Plots

1. **P-value histogram (5.1)** Gives an understanding of the distribution of association p-values. Under the null hypothesis, the p-values of SNPs are expected to be randomly and uniformly distributed from 0 to 1. The histogram is used to assess the significance of the deviation of the p-values from the null. Very strong deviations from a uniform distribution may indicate the presence of an erroneous analysis. You should generate the plot similarly to the previous histogram, but with a range of 0 to 1 and a bin width of 0.05. Below is R code that can be used to generate a p-value histogram.

```
#for these plots, read the results data table with a header
data <- read.table("yourfilename.assoc.linear", header=TRUE, sep = "", fill = TRUE, quote = "")

        hist(as.numeric(data\$P),
        main="p-value Distribution",
        xlab="p-values",
        ylab="Frequency",
        breaks = seq(0,1,0.05),
        freq = TRUE, cex = 3)
```

[8] Please focus on clarity and not length. The reader should have an idea about to perform a similar analysis from reading this file.

2. **Q-Q Plot (5.3)** A quantile-quantile plot plots the observed p-value distribution against the distribution under the null hypothesis. The null hypothesis is that there is no association between the trait being studied and any SNP. We expect the distribution of p-values to be randomly and uniformly distributed. The quantile of a p-value is the proportion of all p-values less than that p-value. In the Q-Q plot, we plot the p-value of each quantile against each other. In other words, we view how significantly the p-value distribution deviates from the null hypothesis, similarly to a histogram. The axes of a Q-Q plot are typically in the $-\log_{10}$ scale, which allows us to visualize the small p-values. The Q-Q plot and histogram are used together to assess how much the observed distribution deviates from the null. We expect to deviate from the null with the smaller p-values, but maintain an overall somewhat uniform distribution. These can be generated for any set of p-values from a PLINK results data table using R:

```
#generates the Q-Q plot
#
#   library("qqman") - calls qqman library
#   data$P      - p-values being plotted
#   main=       - the main title
#
#   https://www.rdocumentation.org/packages/qqman/versions/0.1.2

        library("qqman")
        qq(data$P,
           main = "Q-Q plot"
           )
```

3. **Manhattan Plot (5.4)** This plot visualizes the strength of associations throughout the genome. The SNPs' association p-values are plotted against their positions in the genome. Signals above the standard modern significance threshold ($5.0 \times 10^{-8}$), marked in a red line, and are considered significant. Manhattan plots can also be good indicators for an error in a study. Manhattan plots can be generated using R for a PLINK results data table with:

```
#generates the Manhattan plot
#
#   library("qqman") - loads qqman library
#   data        - the data frame data is being taken from
#   chr="CHR"   - name of column with chromosome number
#   bp="BP"     - name of column with base pair position of SNPs
#   p="P"       - name of column with p-values of SNPs
#   snp="SNP"   - name of column with SNP id
#   main=       - the main title
#
#   https://www.rdocumentation.org/packages/qqman/versions/0.1.2

            library("qqman")
            manhattan(data,
                      chr="CHR",
                      bp="BP",
                      p="P",
                      snp="SNP",
                      main = "Manhattan plot"
                      )
```

4. **Violin Plot (5.5)** A combination of a histogram and box plot. It visualizes the distribution of a trait,

plotted against the genotype of those with the trait. This plot is typically very informative when plotted for the most significant SNPs, as it often demonstrates the presence of a possible allelic effect. That is, when one allele strongly influence the outcome of the phenotype. To generate this plot, find the top SNP in the results using the file containing the top 50 SNPs. Place it in its own text file (for example `top_snp.txt`), and run the following PLINK command.

```
plink --bfile ERAP2_YRI --extract top_snp.txt --recode compound-genotypes --out top_snp_compound_geno
```

Using R, the plot can now be generated with ggplot2. Ensure that the `.ped` file is read and plotted. Since there is no header, the columns are included by numbers in `aes()`.

```
library(ggplot2)
# read the data table with the recoded genotypes
#   header=FALSE - does not read the top row as names for the columns
#   sep = ""     - reads the any length whitespace between text as separator
#   fill = TRUE - matches row length with NA values where applicable
#   quote = ""  - indicates no data is quoted
#
#   https://www.rdocumentation.org/packages/utils/versions/3.6.2/topics/read.table

data = read.table("LRAP_CEU_top_alleles.ped",
                  header = FALSE,
                  sep = "",
                  fill = TRUE,
                  quote = "")

#generates the ggplot
#   data                      - specifies the table being used
#   aes(x=V7, y=V6, fill=V7)  - takes x,y, and fill values from columns
#                               7,6,7 in the data table respectively
#   + geom_violin()           - makes the plot a violin plot

ggplot(
  data,
  aes(x=V7,
      y=V6,
      fill=V7)
  )

+ geom_violin()

#adds features to make plot look better
#   +ggtitle()                - adds main title
#   +xlab()                   - adds x axis title
#   +ylab()                   - adds y axis title
#   +labs(fill=)              - adds a legend
#   +scale_color_manual()     - makes plot a nice colour (don't worry abt this)
#   +theme(legend.position=)  - changes legend position
#   +geom_boxplot(width = 0.1)  - adds a boxplot onto the violin plot to show
#                                 quartiles and median
#
#   http://www.sthda.com/english/wiki/ggplot2-violin-plot-quick-start-guide-r-software-and-data-visualization
```

```
+ ggtitle("gene expression distributions against snp genotypes")
+ xlab("Genotype")
+ ylab("Gene expression")
+ labs(fill = "Genotype")
+ scale_color_manual(values=c("#999999", "#E69F00", "#56B4E9"))
+ theme(legend.position="right" )
+ geom_boxplot(width=0.1)
```

##### 4.2.3. LocusZoom

The data can also be explored through LocusZoom (Boughton *et al.* 2021). This platform visualizes the location of significant SNPs on the genome, shows their relative position to genes, displays linkage disequilibrium using a colour gradient and identifies associations from the GWAS catalogue[9].

1. In order to prepare the data for plotting in LocusZoom, you will need to specify reference alleles within the significant region in your `.assoc.linear` file. This can be done by running the script `ref_allele.R` after modifying it with the appropriate file names.

   **Note:** The following Unix command also creates a LocusZoom format file. To do this on MacOS, you will need to install GNU grep through Homebrew to be able to use the `-w` option (see Section 1.3.6). Change the file names so the `.linear` file is the unsorted results file with no NA values, the `.bim` file is the dataset file, and then the last .txt file will be the name of the output file.

   ```
   paste <(ggrep -w -f <(awk '{print\$2}' YRI_unsorted.linear) YRI_dataset.bim
   | awk 'BEGIN{OFS="\t"}{print\$1,\$4,\$5,\$6}')
   <(awk '{if (NR!=1) {print\$9}}' YRI_unsorted.linear) > LZ_YRI.txt
   ```

2. Once the reference alleles have been specified, upload[10] them to `https://my.locuszoom.org/gwas/upload/`. Enter a descriptive label for the GWAS, specify the build as GRCh37, and attach the file.

3. When the file is uploaded, LocusZoom will open a window which requires you to identify the columns with the chromosome, base pair position and p-value for each SNP. Another parameter which you may optionally add are the reference and alternate alleles. The other parameters may be skipped and the dataset submitted.

4. LocusZoom can display linkage disequilibrium as a colour gradient if the alleles of the top SNP are correctly set to global reference and alternate. Find the top SNP in the sorted results file, and using the SNP ID, locate it in the dbSNP database (Sherry *et al.* 1999):

   `https://www.ncbi.nlm.nih.gov/snp/`

   The alleles will be presented in the format C>A, where C is the reference and A is the alternate allele(s). The reference allele is the one present in the reference sequence of the human genome, and in some cases may be the minor (less common) allele.

[9] The GWAS catalogue is a catalogue of all associations identified in studies.
[10] You may be required to log in. You can use any Google account to do so.

5. Once you have determined the reference and alternative alleles of the top SNP in dbSNP, locate the SNP by the base pair and chromosome in the file uploaded to LocusZoom. (Remember to use the base pair number from the correct genome build). Choose the allele columns such that the reference and alternate alleles match those found on dbSNP. For example, if the reference allele for the top SNP is the C allele, and for that SNP the C allele is in the A2 column of the text document, then A2 should be selected as the reference column and A1 should be selected as the alternate column.

   **Note:** You can also simply try switching A1/A2 when uploading the dataset.

6. After matching reference and alternative allele columns to dbSNP, you may resubmit the data on LocusZoom to display linkage disequilibrium information. LocusZoom will send an email when your plot has been generated, or the page can be refreshed in a few moments.

##### 4.2.4. Questions

- What do you observe in the plots? Is it consistent with what we expect?
- What region of the genome is significant?
- What do you notice about the violin plot compared to the gene expression distribution from section 4.1.1?
- What do you notice from the LocusZoom plots? What genes is the significant SNP located near? What do you notice about the linkage disequilibrium between SNPs around the top SNP (look at the colours)?

#### 4.3. ERAP2 gene expression GWAS on 102 CEU individuals

Now, the goal is for you to independently perform a complete genome-wide association study. Similarly to the YRI individuals, the HAPMAP_3 gene expression dataset (Montgomery *et al.* 2010) contains 104 CEU individuals that overlap with the data cleaned by TCAG (Roslin *et al.* 2016). After examining their principal components, 2 outliers were identified and removed. The GWAS you perform should be analogous to the one in section 4.2, except on these 102 CEU individuals. You may use the above section as a reference.

##### 4.3.1. Questions

- How does the gene expression distribution vary between the YRI and CEU individuals?
- What region of the genome is significant? Are the top SNPs the same between both populations? Is the same region significant?
- Do you observe any differences between the results? Do you expect the results to differ if we perform a GWAS with both populations?
- What do you notice in the LocusZoom plots?

#### 4.4. ERAP2 gene expression GWAS with 88 YRI and 102 CEU individuals

We will now perform the ERAP2 gene expression GWAS including both populations. The individuals in the study are the combined 88 YRI and 102 CEU datasets. The analysis is almost the same as the previous two. However, the differences in population must be accounted for before an association test can take place. This is done by including principal components as covariates, which can be thought of as measures of quantifying population differences. After extracting the data, creating a new binary file and running the MAF and chromosome filters, principal components should be generated in PLINK using

```
plink --bfile ERAP2_YRI_CEU --pca 5 --out ERAP2_YRI_CEU_pc
```

This generates a file with 5 principal components named `ERAP2_YRI_CEU_pc.eigenvec` and their respective magnitudes in `ERAP2_YRI_pc.eigenval`. The process behind PCA is beyond the scope of this workshop. These components should be included in the analysis using

```
plink --bfile ERAP2_YRI_CEU --linear sex hide-covar
--covar LRAP_YRI_CEU_pc.eigenvec --out ERAP2_YRI_CEU_pca_sex
```

Afterwards, you should continue with the usual steps.

##### 4.4.1. Questions

- What do you notice about the gene expression distribution with individuals from both populations?
- What region of the genome is significant? Are the top SNPs the same as before? Is the same region in the genome significant?
- Do you observe any differences between the results?
- What do you notice in the LocusZoom plots?

### 5. Plotting

In this section, the different plots used to interpret GWAS results are discussed in more detail. The scripts to create each plot can be found at `https://github.com/sugolov/GWAS_Workshop`. At the end of each section, a link to the R documentation will be provided.

In practice, you will need to read a data table containing the data you are plotting. In R, this is best done with

```
#reads a data table
#
#   header=TRUE             - reads the top row as names for the columns
#   sep = ""                - reads any length whitespace between text as separator
#   fill = TRUE             - matches row length with NA values where applicable
#   quote = ""              - indicates no data is
#   check.names = false     - does not check package specific naming conventions
#
#   https://www.rdocumentation.org/packages/utils/versions/3.6.2/topics/read.table

results <- read.table("data.txt",
                      header = TRUE,
                      sep = "",
                      fill = TRUE,
                      quote = "",
                      check.names = FALSE)
```

This reads the file with the first line as a header, the separator between cells being any length whitespace character, and fills missing cells so that the dimensions of the dataframe match.

#### 5.1. Histogram

The histogram is a plot that visualizes the distribution of a set of data. In a histogram, the frequency of the occurrence of a particular value is plotted against the value. Below (Figure 3) is a histogram of ERAP2 gene expression in YRI individuals. Note that the gene expression values are continuous, and are separated into bins of width 0.5 to generate the plot.

We see that the gene expression data is bimodal. We can identify the approximate value of the modes, and can see the variance of the data. Other data may be normally or uniformly distributed which is also easy to visualize through a histogram. For example, the p-values of SNPs after a GWAS are expected to be uniformly distributed under the null hypothesis. By making a histogram for these values, we may see if the distribution follows our expectation, i.e. verify that it is close to uniform. However, there are hundreds of thousands of p-values placed in large bins when we generate p-value histograms. For this reason, logarithmically scaled Q-Q plots are used to view what occurs with quantities like p-values, which is discussed in the next section.

To generate a histogram, the `R` commands on the next page can be used.

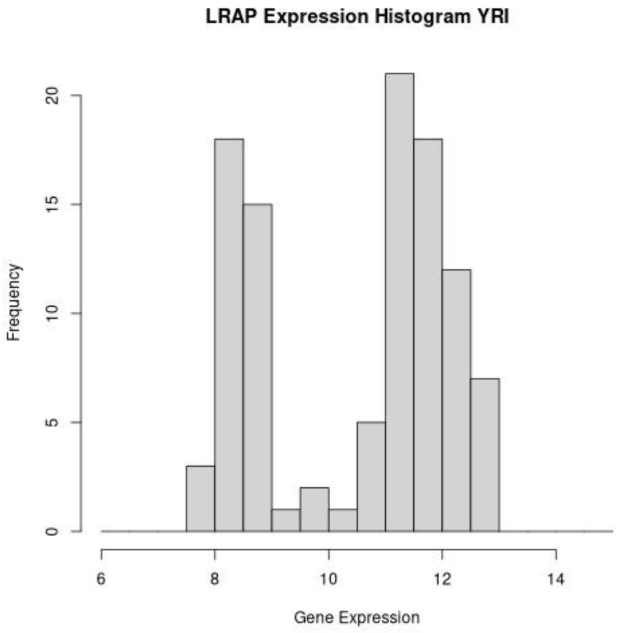

Figure 3: Histogram of ERAP2 gene expression in 91 YRI individuals.

```
#generates the histogram
#
#   as.numeric(data)    - the data being plotted, numeric type of third column in the
#                         data table (gene expression is in the third
#                         column in the phenotype files)
#   main =                - the main title
#   xlab =                - the x axis title
#   ylab =                - the y axis title
#   breaks=seq(a,b,x)  - plots bins of width x taking values from a to b
#   freq = TRUE           - plots frequencies
#   cex = 3               - increases scaling for better resolution

        hist(as.numeric(data),
             main="Gene Expression distribution",
             xlab="Gene Expression value",
             ylab="Frequency",
             breaks = seq(6,14,0.5),
             freq = TRUE, cex = 3 )
```

Note that `breaks = seq(a,b,x)` above creates a histogram with bins from *a* to *b* of size *x* by using the sequence command in R Please see `https://www.rdocumentation.org/packages/graphics/versions/3.6.2/topics/hist` for a more detailed explanation of the hist command.

#### 5.2. Box and whisker plot

The box and whisker plot is a different way to visualize a distribution instead of using a histogram. While the histogram visually emphasizes the data counts present in a range, the boxplot makes explicit the interquartile ranges, mean, median, maximum, minimum, and outliers. Below is an example of a simple boxplot for a normally distributed set of data with mean 0 and standard deviation 1.

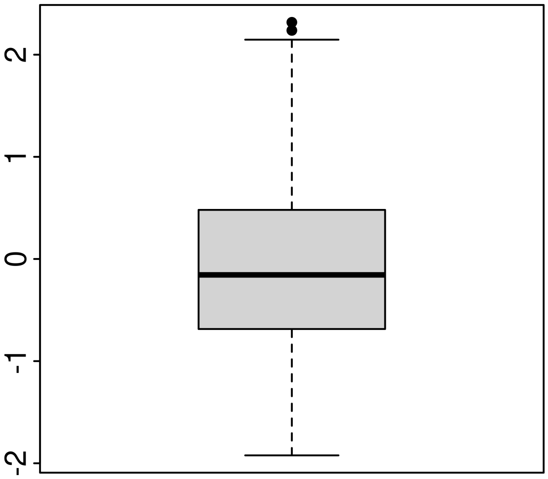

Figure 4: Boxplot of normally distributed data with mean 0 and standard deviation 1.

The above highlights the interquartile ranges of the data. The bottom of the box is the location of the first quartile, the thick line in the middle is the median, and the top of the box is the third quartile. The ends of the whiskers represent the minimum and maximum values in the data set that are not considered outliers. There are two outliers in this data set that are marked as black dots above the maximum. The box and whisker plot emphasizes the variance in the data more than the counts. This plot can be generated in `R` using the `boxplot()` command with

```
#generates normally distributed data set with 50 elements
data = rnorm(50)

boxplot(data)
```

#### 5.3. Q-Q Plot

A quantile-quantile plot plots the observed p-value distribution against the expected distribution under the null hypothesis for some analysis. In a GWAS, the null hypothesis is that there is no association between the trait being studied and any SNP. We therefore expect the distribution of p-values to be randomly uniformly distributed. The quantile of a p-value is the proportion of all p-values less than that p-value. In the Q-Q plot, we plot the quantile of each p-value against each other. In other words, we view how significantly the p-value distribution deviates from the null hypothesis, similarly to a histogram. The axes of a Q-Q plot are typically in the $-\log_{10}$ scale, which causes the plot to focus on the small p-values. The Q-Q plot and histogram are used together to assess how much the observed distribution deviates from the expected distribution under the null hypothesis.

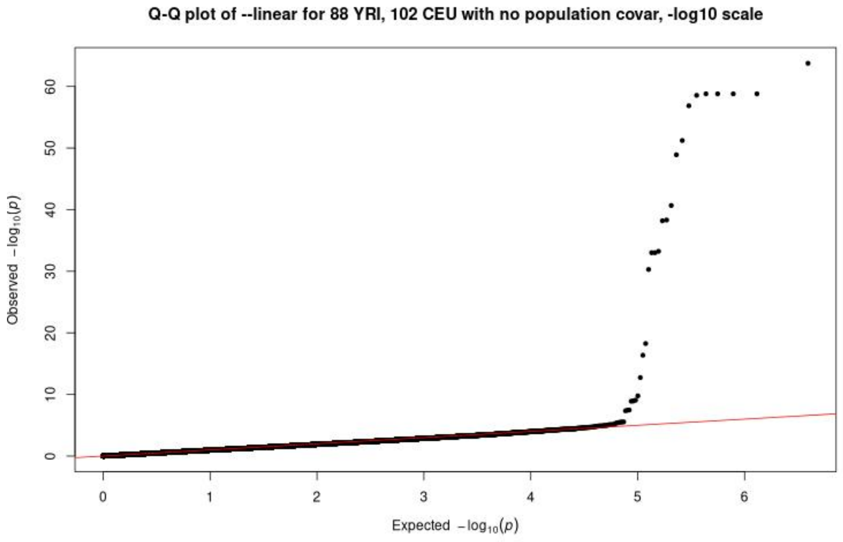

Figure 5: Sample Q-Q plot.

The above Q-Q plot is indicative of results that deviate from the null in some fashion. Note the scaling on the axes: the logarithmic scale causes most of the data points to be clustered near the origin, while the smallest p-values can be examined more closely. Thinking about QQ plots in terms of quantiles is technically correct but also somewhat confusing. It is better to think about a Q-Q plot as a scatter plot with the observed sorted p-values plotted against the expected sorted p-values. To generate a Q-Q plot in R, the following commands may be used. However, without the qqman package, one could generate the same number of random p-values as observed p-values, and plot the sorted lists against each other.

```
#generates the Q-Q plot
#
#   library("qqman") - calls qqman library
#   data\$P      - p-values being plotted
#   main=       - the main title
        library("qqman")
        qq(data\$P,  main = "Q-Q plot" )
# https://www.rdocumentation.org/packages/qqman/versions/0.1.2
```

Please also see the script `linear_plots.R`.

#### 5.4. Manhattan Plot

The Manhattan plot[11] is useful for identifying significant SNPs in the genome. The p-value of each SNP after GWAS is plotted against the location of the SNP in the genome, usually with a $-\log_{10}$ scaling on the p-value. Note, if a Q-Q plot and a Manhattan plot are compared with the same scaling on the observed p-value axis (vertical axis), the p-values should align. That is we can see where the points with the smallest p-value from the Q-Q plot align with their location in the genome in the Manhattan plot. These three plots are therefore essential for understanding GWAS results, as we note the general observed distribution, the deviation from the null of the smallest p-values, and the location of these p-values.

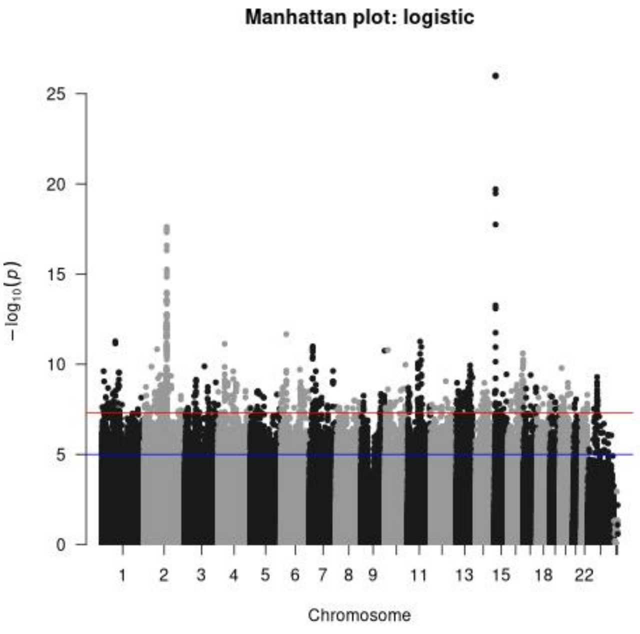

Figure 6: Sample Manhattan plot.

In the above plot we see the chromosomes with strong associations, and the old ($1.0 \times 10^{-5}$) and new ($5.0 \times 10^{-8}$) p-value significance thresholds. The GWAS this plot was generated for was improperly conducted, so you should use the above to get an idea and not as an exemplar.

To generate this plot, you will need to have the `qqman` package installed in R. If you have read a table of PLINK association results, you may run

```
#generates the Manhattan plot
        #
        #   library("qqman") - loads qqman library
        #   data         - the data frame data is being taken from
        #   chr="CHR"    - name of column with chromosome number
        #   bp="BP"      - name of column with base pair position of SNPs
        #   p="P"        - name of column with p-values of SNPs
        #   snp="SNP"    - name of column with SNP id
        #   main=        - the main title
        #
```

[11] When there are multiple associations there are spikes which look like skyscrapers (sort of).

```
#   https://www.rdocumentation.org/packages/qqman/versions/0.1.2

        library("qqman")
        manhattan(data,
                chr="CHR",
                bp="BP",
                p="P",
                snp="SNP",
                main = "Manhattan plot"
                )
```

Please see the script `linear_plots.R`.

#### 5.5. Violin Plot

A violin plot is a combination of a histogram and box and whisker plot. The plots are generated for a single SNP. For the individuals of a given genotype for this SNP, the gene expression distribution is plotted with minimum, maximum, and quartile markers. In other words, the violin plot visualizes the gene expression distribution, plotted against the genotype of the individuals. This plot is typically very informative when plotted for the most significant SNPs, as it can indicate the presence of a strong genetic effect when there are differences in the distribution by genotype.

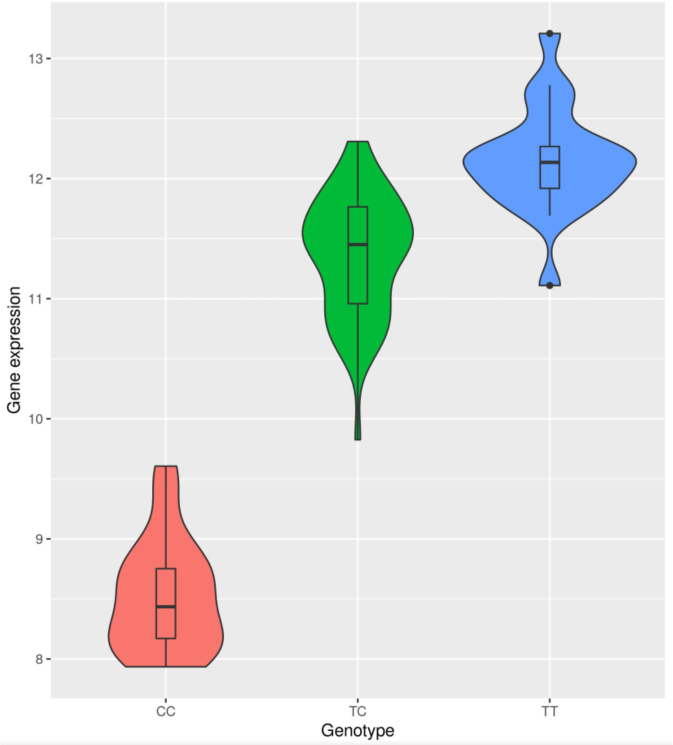

Figure 7: Sample Violin plot.

To generate this plot, find the top SNP in the analysis using the file containing the top 50 SNPs. Place it in its own text file, and run the following PLINK command.

```
plink --bfile binary_file --extract top_snp.txt --recode compound-genotypes --out top_snp_text_file
```

Using R, the plot can now be generated with ggplot2. Ensure that the `.ped` file is read and plotted. Since there is no header, the columns are included by numbers in `aes()`.

```
library(ggplot2)
# read the data table with the recoded genotypes
#   header=FALSE - does not read the top row as names for the columns
#   sep = ""     - reads the any length whitespace between text as separator
#   fill = TRUE - matches row length with NA values where applicable
#   quote = ""  - indicates no data is quoted
#
#   https://www.rdocumentation.org/packages/utils/versions/3.6.2/topics/read.table
```

```
data = read.table("LRAP_CEU_top_alleles.ped",
                  header = FALSE,
                  sep = "",
                  fill = TRUE,
                  quote = "")

#generates the ggplot
#   data                      - specifies the table being used
#   aes(x=V7, y=V6, fill=V7)  - takes x,y, and fill values from columns
#                               7,6,7 in the data table respectively
#   + geom_violin()           - makes the plot a violin plot

ggplot(
  data,
  aes(x=V7,
      y=V6,
      fill=V7)
  )

+ geom_violin()

#adds features to make plot look better
#   +ggtitle()                - adds main title
#   +xlab()                   - adds x axis title
#   +ylab()                   - adds y axis title
#   +labs(fill=)              - adds a legend
#   +scale_color_manual()     - makes plot a nice colour (don't worry abt this)
#   +theme(legend.position=)  - changes legend position
#   +geom_boxplot(width = 0.1)  - adds a boxplot onto the violin plot to show
#                                 quartiles and median
#
#   http://www.sthda.com/english/wiki/ggplot2-violin-plot-quick-start-guide-r-software-and-data-visualization

+ ggtitle("gene expression distributions against snp genotypes")
+ xlab("Genotype")
+ ylab("Gene expression")
+ labs(fill = "Genotype")
+ scale_color_manual(values=c("#999999", "#E69F00", "#56B4E9"))
+ theme(legend.position="right" )
+ geom_boxplot(width=0.1)
```

A good guide about generating violin plots can be found here:

http://www.sthda.com/english/wiki/ggplot2-violin-plot-quick-start-guide-r-software-and-data-visualization

##### 5.6. P-P Plot

The P-P plot is a plot of the p-values of one test against the p-values of another test for a set of variables, in this case SNPs. This plot visualizes whether the p-values are distributed in the same manner, or if different tests yield different statistical significance. A test where the p-values from each test are distributed in the same way will lie on the diagonal, otherwise it will deviate from the diagonal. Since we only perform the `--linear` test, we will not need this type of plot. However, it may be useful if you decide to compare p-values between other tests such as `--assoc` or `--model`. It can also be used to examine p-value differences for analyses including different covariates. It should be noted that Q-Q plots are a type of PP-plot.

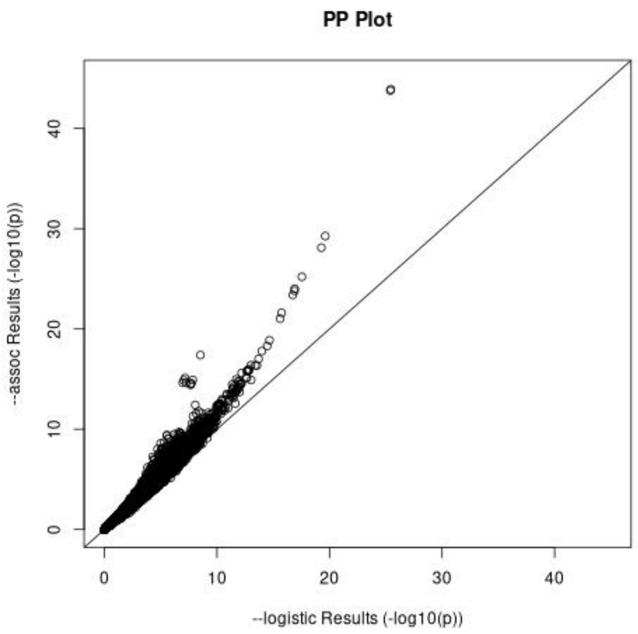

Figure 8: Sample P-P plot.

Due to the logarithmic scaling, most of the data is clustered near the origin. To combat this, hexbin plots are preferable when there is any over-plotting.
After reading the data tables of both tests with the typical PLINK header,

```
xsp <- -log10(as.vector(x_data_table$P))
ysp <- -log10(as.vector(y_data_table$P))
plot(xsp,ysp,main="PP Plot",xlab="test1 Results (-log10(p))",
ylab="test2 Results (-log10(p))",xlim=c(0,45),ylim=c(0,45))+ abline(a=0,b=1)
```

Note that the first two lines scale the axes by $-\log_{10}$. The above uses the default R `plot` command, the documentation to which can be found here:

`https://www.rdocumentation.org/packages/graphics/versions/3.6.2/topics/plot`

##### 5.7. Hexbin Plot

Hexbin plots are a useful way to plot lots of overlapping points, such as the previous P-P plot. Whereas a P-P plot compares each individual p-value for each SNP from two different analyses, hexbin plots show the density of the p-values using a colour gradient. Thus, the same files can be used for a hexbin plot and P-P plot, and the axes are the exact same for the two plots (-log10(p-value)). As mentioned in section 5.6, the plotting of density rather than individual data points is helpful in the case of over-plotting. In Figure 8, the majority of data points are grouped in the same area, which prevents us from interpreting the density of the points. Thus, a hexbin plot would be appropriate in that case.

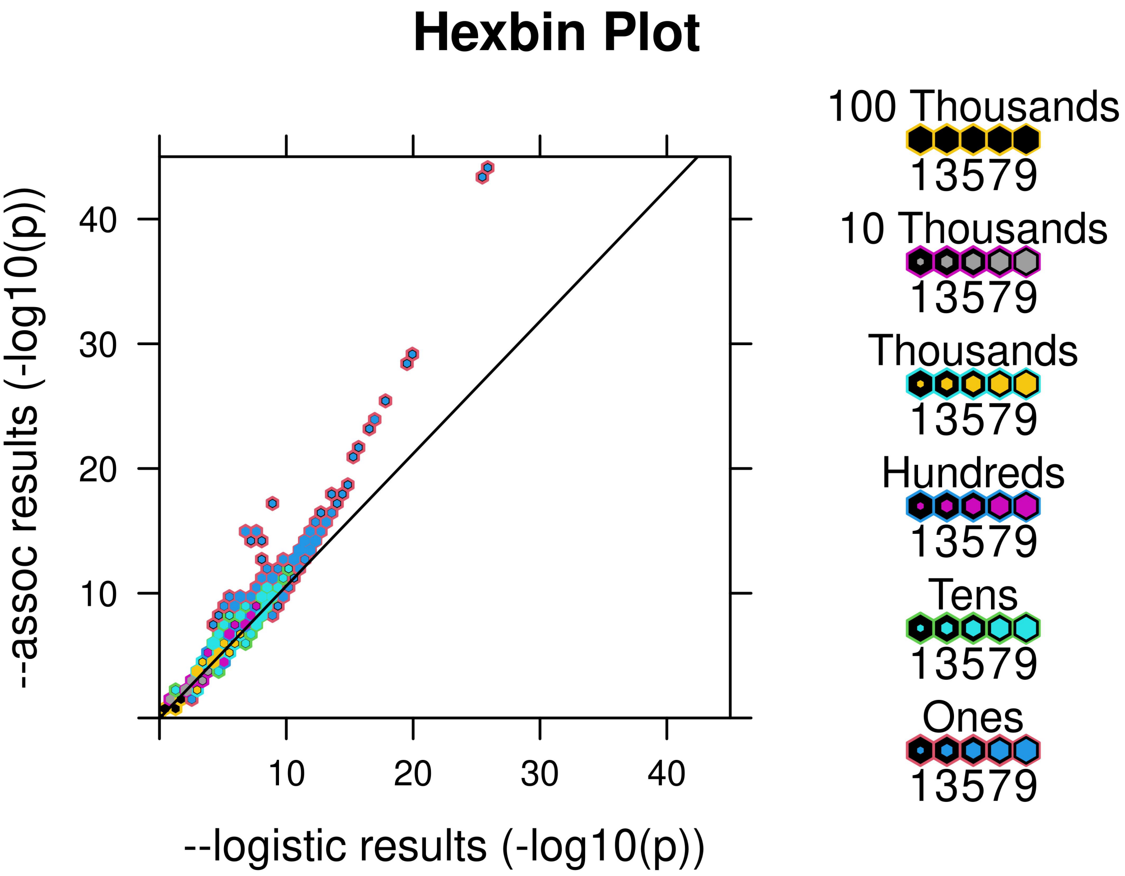

Figure 9: Sample Hexbin plot. This was made using the same data as the P-P plot (Figure 8); please compare the two to understand why a hexbin plot may be more useful than a P-P plot if the data is over-plotted.

Here is sample hexbin plot code.

```
xsp <- -log10(as.vector(xs$P))
ysp <- -log10(as.vector(ys$P))
hexbinplot(ysp~xsp,main="Hexbin Plot",xlab="x axis label",ylab="y axis label",style="nested.lattice"
                        type="r",minarea=0.125,inner=0.1,cex.title=0.75)
hbin <- hexbin(xsp,ysp)
hvp <- hexViewport(hbin)
```

#### 5.8. Scripts

You can find scripts to help in generating these plots on GitHub:

`https://github.com/sugolov/GWAS_Workshop`

### 6. Conclusions

Throughout this workshop, we have found a significant association with the gene expression of ERAP2 and SNPs located in a region of chromosome 5. This reinforces the first part of the central tenet of molecular biology which states that DNA is transcribed to messenger RNA (mRNA). In this workshop, we have shown that the SNPs in this region of chromosome 5 are significantly associated with the amount of mRNA that is transcribed from the ERAP2 gene. The second part of the central tenet of molecular biology is that mRNA encodes proteins which influence the phenotype. This was shown to be true for ERAP2 by Sun *et al.* (2018). It was demonstrated that the effect of SNP rs2927608, within the same region of chromosome 5, on blood protein levels of ERAP2 has a significance of $3.1 \times 10^{-416}$ (ibid.).

Within the GWAS catalog there are numerous studies that have found ERAP2 to be significantly associated with Crohn's disease and general inflammatory bowel disease (IBD) (Franke *et al.* 2010; Jostins *et al.* 2012). Through the LocusZoom plots, the SNPs that were found to be significant in the workshop also lie in a similar genetic region as those discovered in the aforementioned studies.

The SNPs you found have a statistically significant effect on the gene expression of ERAP2. Sun *et al.* (2018) demonstrated that the level of expression of ERAP2 is strongly associated with the level of protein in the blood. Franke *et al.* (2010) and Jostins *et al.* (2012) demonstrated that SNPs located within the region of the ERAP2 gene are significantly associated with Crohn's disease and IBD. In sum, this demonstrates the central tenet of molecular biology: SNPs affect the transcription of the gene to mRNA, which after translation, results in differences in blood protein level between people, and causes a different phenotype to be expressed.

Genome Wide Association studies allow us to understand the genetic risk factors in many diseases to a greater degree than was previously possible. We showed the relative ease with which these studies can be performed after quality control, which demonstrates the potential of GWAS in the future. We hope you will reflect about the significance of these results, and are interested in this powerful area of data science.

### 7. Resources

If you are having difficulties or need clarification, please contact Anton and Eric at respectively.

The PLINK and R documentations are particularly useful for troubleshooting. The PLINK v1.90 documentation should be checked first and most trusted, but the PLINK v1.07 (old) documentation is sometimes referenced or useful. Please refer to these links first if PLINK gives you any errors: the website covers the correct input for a given command. If the errors persist, please contact either Anton and/or Eric. Outside of this workshop, it is likely that there is an error with the formatting of your data; the next step in trouble shooting would be to check that all of the files are properly formatted.

For R, the syntax and parameters of any command are available at the official R documentation or the documentation for the specific package (such as `qqman`). StackExchange posts are also useful, as most introductory questions are answered well. The provided R scripts will have some form of annotation about how they work. If something is unclear, please do ask.

For LocusZoom, the attached FAQ may help in case your plots are not being generated due to errors, or if the colour for linkage disequilibrium is not being shown. The dbSNP database has also been linked here for easy access. The home page of the dbSNP database has a brief set of instructions on how you can search it, and links to a more detailed set of instructions you can use to narrow down your search results.

Lastly, we have provided one LocusZoom link for each of the three analyses you should have performed. Please compare these results to your own to check that you have successfully performed all steps as outlined in Sections 4.2, 4.3 and 4.4.

- **PLINK v1.07 Documentation:** `https://zzz.bwh.harvard.edu/plink/`
- **PLINK v1.90 Documentation:** `https://www.cog-genomics.org/plink2/`
- **R Documentation [12]:** `https://www.rdocumentation.org/`
- **LocusZoom FAQ:** `https://my.locuszoom.org/about/`
- **dbSNP Home Page:** `https://www.ncbi.nlm.nih.gov/snp/`
- **LocusZoom YRI Analysis Results:** `https://my.locuszoom.org/gwas/841221/`
- **LocusZoom CEU Analysis Results:** `https://my.locuszoom.org/gwas/191134/`
- **LocusZoom YRI and CEU Analysis Results:** `https://my.locuszoom.org/gwas/345564/`

[12] It is better to search specific problems, but this is good for syntax.

We are greatly thankful for the efforts of Dr. Sun and Dr. Paterson in teaching us how to perform GWAS and advising this manual.
